# Insulin-like Growth Factor-1 Regulates the Mechanosensitivity of Chondrocytes by Modulating TRPV4

**DOI:** 10.1101/2020.03.10.985713

**Authors:** Nicholas Trompeter, Joseph D. Gardinier, Victor DeBarros, Mary Boggs, Vimal Gangadharan, William J. Cain, Lauren Hurd, Randall L. Duncan

**Affiliations:** Biomedical Engineering, University of Delaware, Newark, DE; Biomechanics and Movement Science; Department of Biological Sciences; Bone and Joint Center, Henry Ford Hospital, Detroit, MI; Department of Biology, University of Michigan-Flint, Flint, MI

**Keywords:** Chondrocytes, Mechanosensitivity, IGF-1, TRPV4 channels, Intracellular Calcium, Hypotonic Swelling, ATP release

## Abstract

Both mechanical and IGF-1 stimulation are required for normal articular cartilage development and maintenance of the extracellular matrix. While much effort has been made to define the signaling pathways associated with these anabolic stimuli, we focused on how these pathways interact to regulate chondrocyte function. The Transient Receptor Potential Vanilloid 4 (TRPV4) channel is central to chondrocyte mechanotransduction and regulation of cartilage homeostasis. However, the mechanism by which TRPV4 is mechanically gated or regulated is not clear. In this study we propose that insulin-like growth factor 1 (IGF-1), which is important in regulating matrix production during mechanical load, modulates TRPV4 channel activity. Our studies indicate that IGF-1 reduces hypotonic-induced TRPV4 currents, and intracellular calcium flux by increasing stress fiber formation and apparent cell stiffness. Disruption of F-actin following IFG-1 treatment results in the return of the intracellular calcium response to hypotonic swelling. Furthermore, we highlight that IGF-1 suppresses TRPV4 mediated calcium flux through the MAP7 binding domain (aa. 798-809), where actin binds to the TRPV4 channel. IGF-1 treatment differentially influences the intracellular calcium flux of HEK 293 cells stably expressing either wild-type or mutant (P799L or G800D) TRPV4 during hypotonic challenge. A key down-stream response to mechanical stimulation of chondrocytes is ATP release. Data here indicate that activation of TRPV4 through hypotonic swelling induces ATP release, but this release is greatly reduced with IGF-1 treatment. Taken together this study indicates that IGF-1 modulates TRPV4 channel response to mechanical stimulation by increasing cell stiffness. As chondrocyte response to mechanical stimulation is greatly altered during OA progression, IGF-1 presents as a promising candidate for prevention and treatment of articular cartilage damage.

## INTRODUCTION

Osteoarthritis (OA) is the leading cause of disability among American adults (CDC, 2010; Helmick et al, 2008) and is characterized by the degradation of the articular cartilage that provides load support and smooth articulation for diarthrodial joints. Articular cartilage homeostasis requires a continuous balance between repair and degradation (Carter et al., 2004; Trippel, 1995). Chondrocytes embedded within the extracellular matrix (ECM) of cartilage contribute to this remodeling process by responding to biomechanical and biochemical signals to synthesize or breakdown the ECM (Mariani et al., 2014). The mechanical environment experienced by chondrocytes can initiate both catabolic and anabolic activity (De Croos et al., 2006; Nebuleng et al., 2012) depending on the magnitude and type of stimuli (Kim et al,. 1994; Mawatari et al., 2009). These actions are mediated, in part, by signaling pathways that include growth factors and receptors (Neu et al., 2007; Vincent et al., 2007) mitogen-activated protein (MAP) kinases (Fanning et al., 2003), Rho guanosine-5’-triphosphate (GTP)ases (Haudenschild et al., 2008), nitric oxide (Mawatari et al., 2009; Oh et al., 2003), integrins (Lee et al., 2002) and ion channels (Lee et al. 2000). Among these agents, insulin-like growth factor-1 (IGF-1) is unusual in that it stimulates both mitogenic and anabolic functions and inhibits catabolic activity in articular chondrocytes (Madry et al., 2001; Novakofski et al., 2009; Sah et al., 1991; Tyler, 1989). These diverse actions are mediated by a complex interplay among multiple signaling pathways including phosphoinositide-3-kinase (PI3K), MAP kinases and Rho GTPases (Fanning et al., 2003; Fortier et al., 2004; Oh et al., 2003). Mechanical forces and IGF-1 interact to regulate articular chondrocytes (Bonassar et al., 2001; Bonassar et al., 2000; Jin et al., 2003), however the mechanisms responsible for this interaction between a biochemical stimulus and a biomechanical stimulus remain unclear.

Calcium (Ca^2+^) signaling is essential to chondrocyte function, regulating many of the genes associated with anabolic activities (Matta et al., 2013). Transient Receptor Potential (TRP) channels are a superfamily of cation-selective channels that modulate calcium signaling through Ca^2+^ influx and modulating Ca^2+^ release and are expressed in almost every tissue and cell type in humans (Pedersen et al., 2005). The TRPV4 channel is central to this Ca^2+^ signaling in chondrocytes and has been shown to regulate SOX9 expression (Muramatsu et al. 2007), a key transcription factor involved in matrix synthesis. TRPV4 channels are gated by a wide range of physical and chemical stimuli including Ca^2+^, temperature, arachidonic acid metabolites, phorbol esters and some synthetic and natural agonists (for review, see (Nilius et al., 2003). TRPV4 has been shown to be involved in the cellular response to changes in osmolarity (Liedtke, 2006; Nilius et al., 2001; Nilius et al., 2003) and mechanical stimulation (Flockerzi et al., 2007; Gao et al., 2003; Loukin et al., 2010; Mizoguchi et al., 2008), both of which are integral to normal cartilage function. The importance of the TRPV4 channel to cartilage homeostasis is illustrated by targeted deletion of TRPV4 in mice that results in severe joint damage resembling osteoarthritis. Chondrocytes isolated from these mice exhibited a loss of the chondrocyte response to osmotic swelling, suggesting a role for the TRPV4 channel in maintenance of cartilage structure (Clark et al., 2010). However, the mechanism through which mechanical stimulation gates the TRPV4 channel, as well as the role of this channel in mechanotransduction is unclear.

Chondrocyte stiffness has been shown to be significantly greater in chondrocytes isolated from OA cartilage than those isolated from normal articular cartilage (Trickey et al., 2004). However, it remains unclear what causes the changes in cell stiffness and how such changes influence cellular responses to mechanical stimuli. We have previously shown that osteoblasts subjected to FSS increase the polymerization of F-actin through activation of RhoA GTPase that, in turn, increases the stiffness of the cell (Gardinier et al., 2014). Previous studies have shown that IGF-1 treatment of primary chondrocytes also increases cell stiffness and F-actin formation (Leipzig et al., 2006) and that this increase in F-actin by IGF-1 treatment results from activation of a Rho GTPase (Novakofski et al., 2009). We have also shown that actin organization regulates the activity of the mechanosensitive channel found in these osteoblasts (Zhang et al., 2006). Suzuki et al. also suggest that the TRPV4 channel is regulated by the actin cytoskeleton through binding of F-actin to the Microtubule Associated Protein Binding 7 (MAP7: amino acids 798-809) domain of TRPV4 (Suzuki et al., 2003). Two point mutations within this domain (P799L and G800D) results in metatropic dysplasia, a severe skeletal dysplasia, in children (Hurd et al., 2015). We have recently shown that these mutations cause a gain of function of the TRPV4 channel that alters chondrogenesis and chondrocyte intracellular calcium influx in human chondrocytes (Hurd et al., 2015). In this study, we have expressed these mutations in HEK 293 cells, a human kidney cell line that do not normally express TRPV4 channels, to determine how actin cytoskeletal organization regulates TRPV4 gating. We postulated that the TRPV4 channel was essential to Ca^2+^ signaling and mechanotransduction in chondrocytes and that IGF-1 would modulate the activity of this channel through alteration in the actin cytoskeletal organization.

## METHODS

### Cell Culture

ATDC5 mouse chondrocyte-like cells (Sigma-Aldrich, St. Louis, MO) were cultured in normal medium consisting of Dulbecco’s Modified Eagle Medium (DMEM, Sigma), 10% FBS (Gibco, New York, NY), 100 U/ml penicillin G, and 100 μg/ml streptomycin (P/S, Mediatech, Manassas, VA) and buffered to pH 7.35 with 26 mM NaHCO_3_ (Fisher Scientific, Pittsburgh, PA). HEK 293 cells transfected with either wild-type TRPV4 (WT), with a proline to leucine mutation at the 799 amino acid site of TRPV4, or a glycine to aspartate mutation at the 800 amino acid site of TRPV4 were grown in Minimal Essential Media (MEM, Corning) supplemented with 10% FBS and 1% P/S. Cells were maintained in a humidified incubator at 37°C with 5% CO_2_/95% air and passaged once cells reached 75% confluence. Once cells reached 80-90% confluence, they were serum starved for 16 hours in reduced serum media (0.2% FBS, 1% P/S).

### Generation of stably expressing HEK TRPV4 cell lines

To create HEK 293 cells that express both the wild type and one of the mutant proteins in a 1:1 ratio, we used two strategies. First, Gibson assembly was used to create a plasmid that produces a tricistronic mRNA coding for the WT TRPV4, either P799L or G800D, and daGFP (referred to as tricistronic). Cells successfully transfected with this construct fluoresce green and be heterozygous for TRPV4. In the second strategy, Gibson assembly was used to clone the WT TRPV4 cDNA into pSF-CMV-EMCV-Neo (Oxford Genetics). This construct (referred to as bicistronic-Neo) is similar to bicistronic GFP. We used this plasmid to co-transfect HEK cells along with a bicistronic-GFP construct expressing one of the mutations. HEK 293 cells were seeded at 8,000-10,000 cells/cm^2^ on type 1 collagen (10 μg/cm^2^, Sigma-Aldrich) coated 35 mm polystyrene dishes (Falcon) and were allowed to grow overnight in MEM with 10% FBS and 1% P/S. Transfection was accomplished using MEM media containing 200 μl of transfection mixture, 4 μL of Lipofectin (Life Technologies) and 1 μg of plasmid. Cells were incubated for 24 hr at 37 in 5% CO_2_/95% air. Following this incubation, the transfection mixture was replaced with growth media and were grown for an additional 48 hr before being replaced with growth media supplemented with 200 μg/ml G418 sulfate (Gibco).

Cells were allowed to grow for an additional 48 hours before being plated as single cells in a 96-well plate (Corning) and checked for expression of GFP under a microscope. Colonies expressing GFP were grown for 4 days before transferring to T25 flasks (Corning) and grown until 70% confluent. Any cells frozen where thawed, seeded onto tissue culture flasks and subsequently checked for GFP expression.

### Electrophysiological recordings

ATDC5 cells were cultured in DMEM with 10% FBS and 100 U/ml penicillin G, and 100 μg/ml streptomycin on 15mm round coverglass (Ted Pella, Inc., Redding, CA). Pipettes were manufactured by pulling 100 μL capillary tubes (VWR, Arlington Heights, IL) to a tip diameter of ~0.5 micron (tip resistance = 1-5 MΩ) using a dual-stage pipette puller (Narishige, East Meadow, NY). Pipettes were flame polished and coated with wax to reduce capacitance artifacts. Cells were bathed in an extracellular-like solution (ECF) composed of (in mM): NaCl 150, CsCl 6, MgCl_2_ 1, CaCl_2_ 5, HEPES 10 and glucose 10, titrated to pH 7.4 with CsOH (total osmolarity 320mOsm). Pipettes were filled with an intracellular-like solution (ICF) containing (in mM): CsCl 20, CsAspartate 100, MgCl_2_ 1, HEPES 10, Mg_2_ATP 4, CaCl_2_ 0.08 and BAPTA 10, titrated to pH 7.4 with CsOH. Whole cell recordings were performed with an Axopatch 200B amplifier (Axon Instruments, USA) equipped with a Digidata 1332A A/D converter. Recordings were performed using a ramp protocol that clamps the cell at 0mV, then ramps the membrane voltage from −80mV to +80mV over 1000ms. The cell then is returned to 0mV for 12.5s prior to restarting the protocol. Data were collected on an IBM compatible PC using pClamp version 10.0 software (Axon Instruments).

To determine the effect of hypotonic swelling on TRPV4 whole cell currents, the ECF was diluted to approximately 110 mOsm (75% change in osmolarity), perfused into the patch chamber and changes in whole cell current recorded using the same ramp protocol as described above. To determine if the TRPV4 channel was responsible for the currents induced by hypotonic swelling, cells were pre-incubated with 10μM RN-1734 (Sigma), a specific inhibitor for TRPV4 channels, for 15 minutes prior to swelling. To determine the effects of IGF-1 on TRPV4 channel activity, the cell medium was replaced with DMEM containing 0.2% FBS and 300ng/mL IGF-1 (human recombinant, Sigma) for 3 hours. Electrophysiology measurements were then performed to determine if IGF-1 pretreatment altered TRPV4 currents induced by hypotonic swelling. All data were analyzed using Clampfit version 10.0 software (Axon Instruments).

### Mechanical Stimuli and Pharmacological Treatments

ATDC5 cells were subjected to the mechanical stimuli of hypotonic swelling (HS) by adding equal volume of water. The tension generated along the cell membrane mimics that experienced by chondrocytes under dynamic loading of cartilage tissue. Changes in the mechanosensitivity of ATDC5 cells to HS was examined when cells were pre-treated with 300 ng/ml IGF-1 for 3 hours prior to HT, or pre-treated for 10 min with the general TRPV channel inhibitor, ruthenium red (10 μM),, or pre-treatment for 10 min with the TRPV4-specific inhibitor, RN-1734. The concentrations of IGF-1, ruthenium red, and RN-1734 were maintained during HS. In order to determine the role of cytoskeleton organization on mechanosensitivity following IGF-1 treatment, cells were treated with 1 μM cytochalasin D for 30 minutes prior to HS in both IGF-1 treated and un-treated cells.

### Immunocytochemistry and confocal microscopy

To image the actin cytoskeleton following various treatments, cells were first washed with PBS for 5 minutes and then fixed with 4% paraformaldehyde (Electron Microscopy Sciences, PA, USA) in PBS containing 0.1% Triton X-100 (Sigma) for 30 min on ice. Cells were then washed with PBS and incubated with blocking buffer containing 3% BSA (Sigma) at room temperature for 1 hour. After blocking, cells were incubated with Alexa Fluor^®^ 488 phalloidin (1:1000, Life Technologies, Grand Island, NY) to label F-actin green. Slides were then mounted using the SlowFade^®^ Antifade kit (Life Technologies). Fluorescent images were obtained using a Zeiss LSM710 laser scanning confocal microscope with a 63X oil immersion lens at the midplane of the cells.

### Atomic Force Microscopy

The apparent stiffness of individual cells was measured using an Atomic Force Microscope (AFM: BioScope II, Veeco Inc., Plainview, NY) mounted on an inverted optical microscope. Soft microlever probes (MLCT-AUNM, Veeco Inc.) with a conical tip and a spring constant of 0.01 N/m were first calibrated using a thermal fluctuation method in fluid. Individual cells with normal morphology were then identified under the optical microscope and positioned under the probe. An area of 30 μm × 30 μm was first scanned at a speed of ~3 μm/sec to generate a topographic map of an individual cell. Seven to ten points were then selected over the cell body at the nuclear and peri-nuclear regions for indentation measurements. The peripheral region of the cell was avoided due to its relatively thin cell height and influence of the rigid substrate. At each selected point over the cell body, the cantilever tip indented the cell membrane at a speed of ~2.5 μm/sec until a force of 100 pN was reached, which typically resulted in an indentation depth of < 70 nm (10% of the cell height). The elastic modulus at each point was estimated from the recorded force-deflection curve using a Hertz based model [6] as defined by the following:

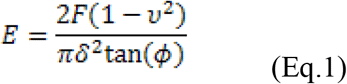

where E is the apparent stiffness, F is the cantilever force measured by the AFM, υ is the Poisson ratio of the cytoplasm (υ = 0.4) (Mak et al., 1987), φ is the opening angle of the conical cantilever tip (φ = 35°). The indentation depth (δ) was calculated by subtracting the cantilever deflection from the piezo displacement of the probe. The apparent stiffness for a given cell was defined by the average elastic modulus across each point measured. A total of 5 to 6 cells were measured following a given treatment, and each experiment was then repeated 3 times. The average cell stiffness for each treatment was compared to static controls and reported as a fold increase. Static controls were not subjected to any form of treatment or mechanical stimuli.

### Calcium Imaging

To measure the intracellular calcium concentration ([Ca^2+^]_i_) of ATDC5 cells in response to HS, cells were first rinsed with Hanks’ balanced saline solution (HBSS, Sigma) and then loaded with fluorescent Ca^2+^ probes (3 μM of fura-2 AM or 5.69 μM of Fluo-4 AM, Life Technologies) in HBSS for 30 minutes at 37°C. The cells were again rinsed with HBSS and incubated at 37°C for 15 minutes in HBSS alone. During mechanical loading, changes in [Ca^2+^]_i_ were recorded using a ratiometric video-image analysis apparatus (Intracellular Imaging, Cincinnati, OH, USA) on a Nikon inverted microscope with a Nikon 30× fluor objective when loading the cells with fura-2 AM. The ratio of emitted light at 340 and 380 nm excitation was determined (F340/F380) from consecutive frames and used to determine the [Ca^2+^]_i_ of selected cells based on previous calibration of known standards. When loading cells with Fluo-4 AM, measurements were taken on a Zeiss 5 Live DUO Highspeed Confocal with a 10×/0.3A water immersion objective. Zen imaging software was used to collect relative fluorescence every 500ms. Cell fluorescence was measured using the region of interest tool to select cells with a circular ROI set to 25 pixels. Zen software compiles fluorescence measurements for each cell in a table, which was then exported to excel to allow for peak over baseline analysis. A basal level of [Ca^2+^]_i_ was first recorded for 1 minute, after which HS was applied by adding equal volume of water to the cells immersed in HBSS. The influence of IGF-1 on the [Ca^2+^]_i_ response was determined by treating either ATDC5 or HEK TRPV4 WT and TRPV4 mutant cells with 300 ng/ml of IGF-1 for 3 hours prior to loading with fura-2AM or Fluo-4AM. For dose response experiments with IGF-1, 0, 1, 100, and 300 ng/ml were added to ATDC5 cells for 3 hr before loading of cells. Time course studies of 300 ng/ml IGF-1 treatment on ATDC5 cells were also completed with pretreatment times of 0, 30 minutes, 60 minutes, 120 minutes, and 180 minutes utilized prior to loading with fura-2 AM. Cytoskeleton organization was also manipulated by treating ATDC5 cells with 5μM of cytochalasin D for 30 minutes prior to HS.

### ATP Assay

Purinergic signaling in response to mechanical loading has a large influence on the metabolism of chondrocytes. The release of ATP was measured from media samples extracted 5 minutes after the onset of HS to chondrocytes. The ATP concentration of each sample was measured using a bioluminescence assay kit (ATP Bioluminescence Assay kit HTS II; Roche, Indianapolis, IN). Light emitted as a result of the reaction between D-luciferin and luciferase was detected using a 96 well micro-injector plate reader (POLARstar OPTIMA, BMG LABTECH GmBH). To normalize the measured ATP release to cell protein, the ATDC5 cells were washed with PBS immediately after HS, and then incubated at −20°C with lysis buffer (5 mM HEPES, 150 mM NaCl, 26% glycerol, 1.5 mM MgCl_2_, 0.2 mM EDTA, 0.5 mM dithiothreitol, and 0.5 mM phenylmethylsulfonyl fluoride, and protease cocktail inhibitor (Sigma;10.4 mM 4-(2-Aminoethyl) benzenesulfonyl fluoride hydrochloride, 8 μM aprotinin, 400 μM bestatin, 140 μM E-64, 200 μM leupeptin, and 150 μM pepstatin A). The protein samples were stored at −80°C for further analysis. The protein concentration of each sample was determined using a BCA assay (BCA Protein Assay Kit, Pierce, Rockford, IL) and the 96-well micro-injector plate reader.

### Statistical Analysis

The change in cell stiffness and ATP release was reported as the average ± SD fold increase relative to static controls unless otherwise stated. Significance between groups was determined using one-way or two-way analysis of variance (ANOVA) with a Tukey posthoc test to determine significance when multiple comparisons in the study were made. Significance was defined by a *p-value* < 0.05. Data for a given outcome measurement was pooled together in the case that no significant difference was found for a given treatment between different passages of cells.

## RESULTS

### TRPV4 channel activity is suppressed in the presence of IGF-1

To determine the effects of IGF-1 on TRPV4 currents in ATDC5 cells, differences in whole-cell currents were determined when the patch clamp voltage was ramped from −80 mV to +80 mV in ATDC5 cells that were treated with either IGF-1 or RN1734 followed by hypotonic swelling. Representative traces are shown in Fig. 1A. Untreated, non-swelled (static) cells exhibited basal currents of 45.4pA/pF ± 1.6 over the voltage clamp range (n=13). When these cells were hypotonically swelled, whole cell currents increased 4-fold in comparison to static controls (254.5pA/pF ± 8.5; n=5). Pre-treatment with 300ng/ml IGF-1for three hours prior to patch completely blocked the increase in currents during hypotonic swelling, resulting in currents comparable to those seen in static cells (34.1pA/pF ± 1.7; n=8). Pretreatment with the TRPV4-specific antagonist, RN1734, prior to hypotonic swelling blocked whole cell currents in hypotonically swelled ATDC5 cells (33.8pA/pF ± 2.9; n=4). These data indicate that the swelling activated currents are mediated by activation of the TRPV4 channel.

**Figure 1:**
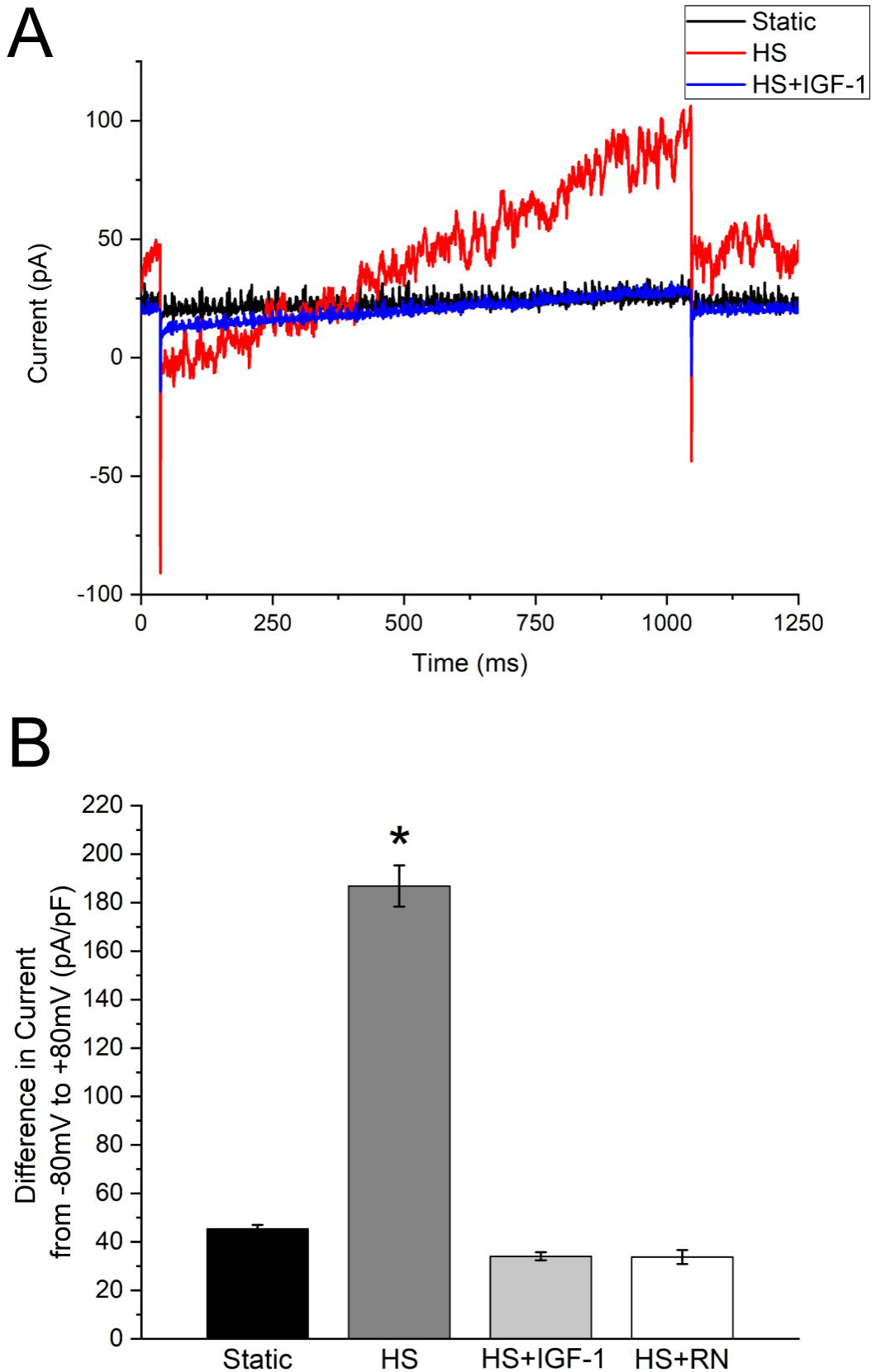
TRPV4 channel activity is altered in the presence of IGF-1. **A]** Representative whole cell current traces from ATDC5 cells that underwent hypotonic swelling (HS) after treatment with the IGF-1 (300ng/mL) for 3 hours, RN1734 (10μM) for 15 minutes, cells that were left untreated, and static controls. Current was greatly reduced when cells were pretreated with IGF-1. **B]** Compilation of average difference in response from −80mV to +80mV. Hypotonic swelling of cells resulted in a 4-fold increase in current in comparison to static control. Hypotonic swelling had no effect on cells that were pretreated with IGF-1. Inhibition with the TRPV4 blocker, RN1734, followed by hypotonic swelling resulted in complete reduction in current. (Tukey-Kramer; * indicates p-value is < 0.05 in comparison to all other conditions)

### Actin Stress Fiber Formation and Cell Stiffness

The cytoskeleton is known to contribute to the mechanical behavior of chondrocytes, specifically cell stiffness (Leipzig et al., 2006), which can greatly influence mechanotransduction. IGF-1 increased actin stress fiber formation (ASFF) in ATDC5 cells as early as 2 hours after treatment (Fig. 2A). In addition, IGF-1 changed the ATDC5 cell morphology to a flatter, elongated shape with stellate processes. ASFF appeared to peak after 3 hours treatment with IGF-1 and was associated with ~ 4-fold (12 kPa) increase in cell stiffness compared to static untreated controls (3.5 kPa) (Fig. 2B). The increase in cell stiffness due to IGF-1 was dependent on F-actin based on the decrease in cell stiffness following cytochalasin D treatment.

**Figure 2:**
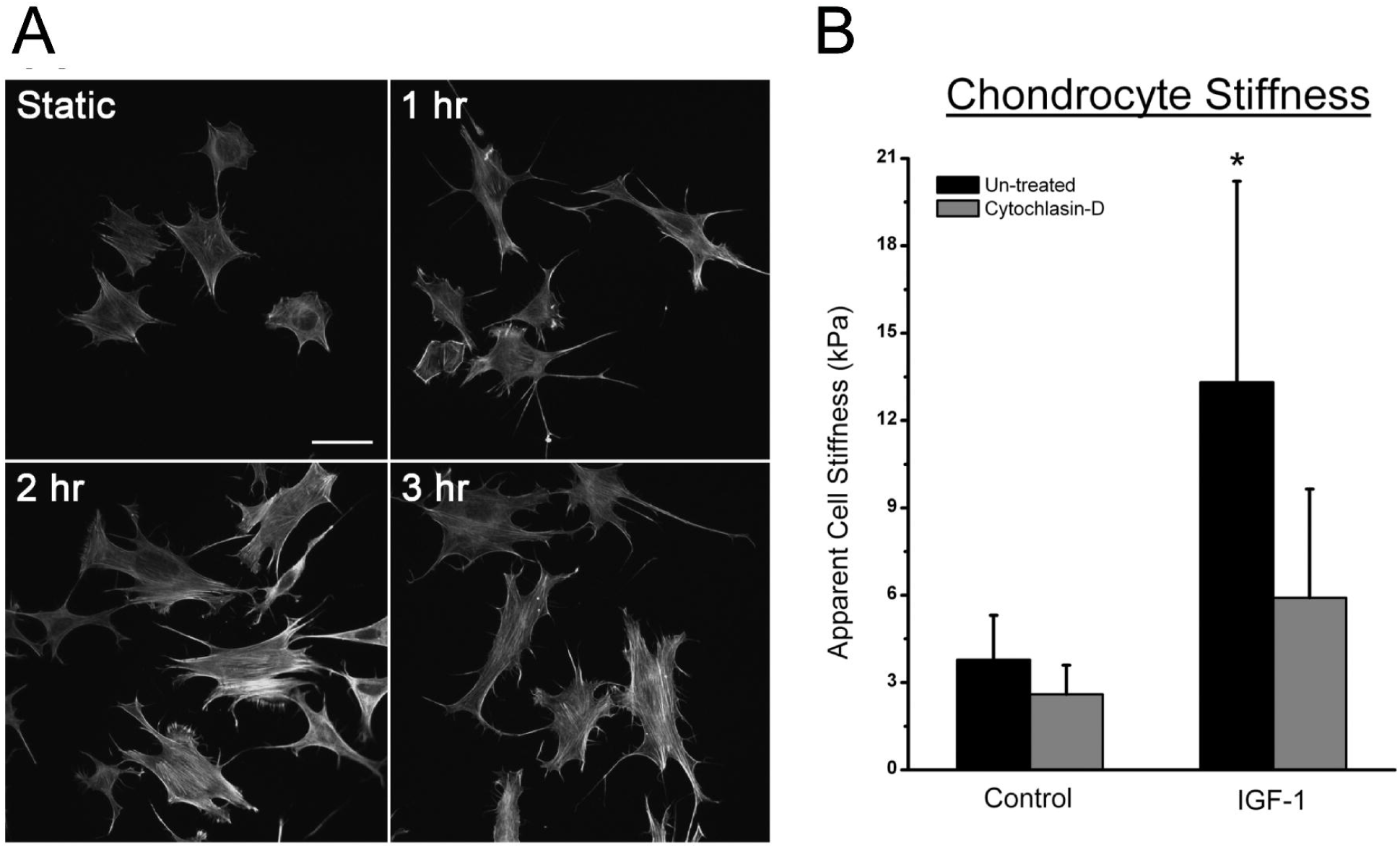
IGF-1 increases actin stress fiber formation and cell stiffness among ATDC5 cells. **A]** Time course of IGF-1 treatment indicates an increased stress fiber formation and cell spreading after 1 hour, with the peak occurring 3 hours. Bar indicates 50 μm. **B] A**pparent cell stiffness of ATDC5 cells was significantly higher after 3 hour treatment with IGF-1. The increased stiffness due to IGF-1 was dependent on the F-actin organization based on the decreased in cell stiffness following cytochlasin D. (* indicates p-value is < 0.05 compared to controls)

### TRPV4 Channel Mediated Calcium Response

One of the earliest responses to mechanical loading in chondrocytes is an influx of calcium into the cell that, in turn, initiates signaling cascades through wide range of different intracellular pathways (Matta et al., 2013). When subjecting ATDC5 cells to HS, we a found a significant increase in the number of cells exhibiting a calcium response, defined as a 50% increase in [Ca^2+^]_i_ above baseline, as well as a significant increase in the magnitude of the [Ca^2+^]_i_ response (Fig. 3). To determine if TRPV channels mediate the [Ca^2+^]_i_ response to HS, ATDC5 cells were pre-treated with the general TRPV inhibitor, ruthenium red. This non-specific inhibition of TRPV channels significantly decreased both the number of cells that responded to HS and the magnitude of the [Ca^2+^]_i_ response to HS (Fig 3). Addition of the TRPV4-specific inhibitor, RN1734, attenuated the number of cells responding and magnitude of [Ca^2+^]_i_ response to HS to the same degree as ruthenium red.

**Figure 3:**
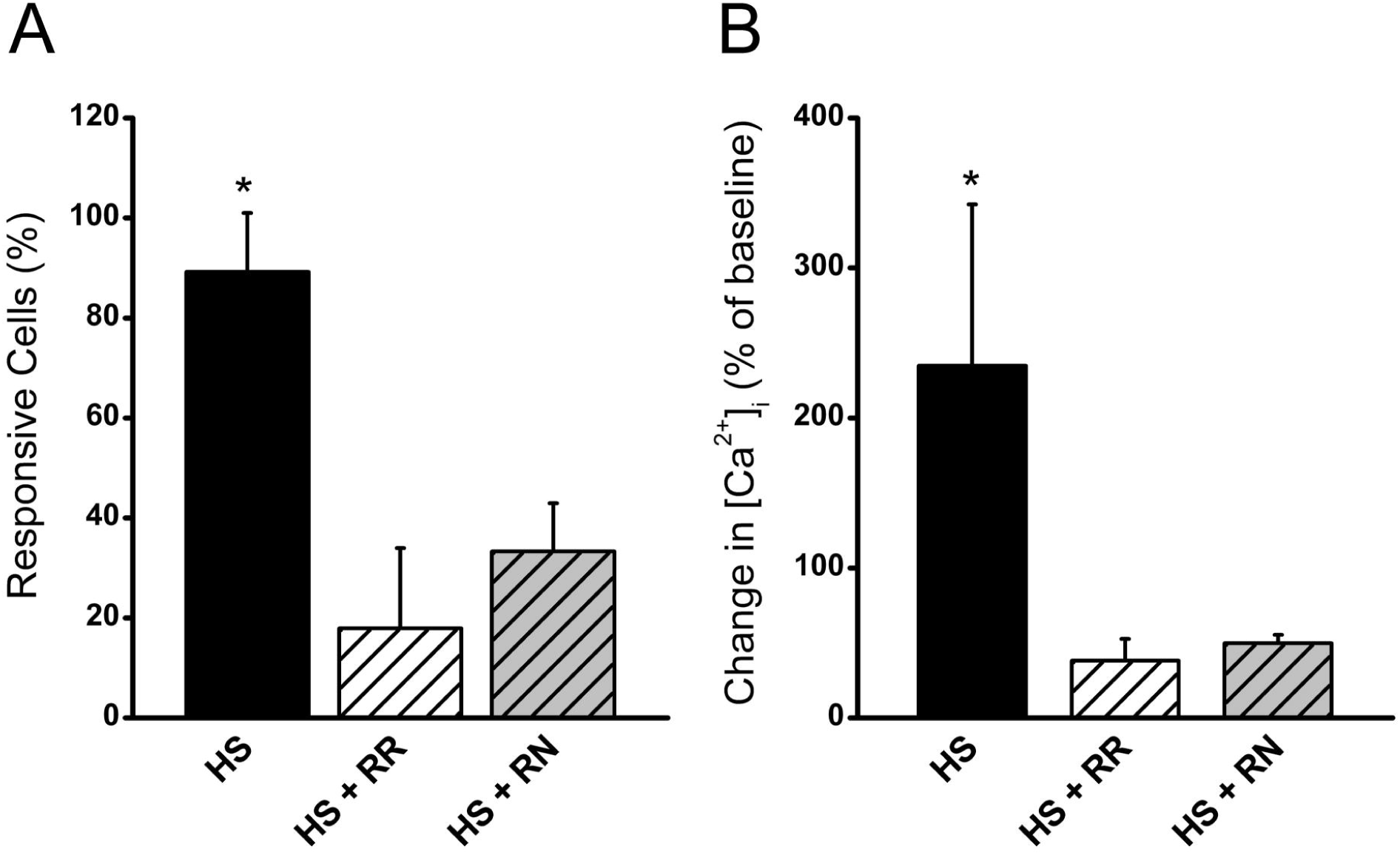
The [Ca^2+^]_i_ response to HS is mediated through TRPV4 channel. **A]** The number of cells responding and **B]** the change in [Ca^2+^]_i_ as a percentage of baseline when ATDC5 cells were treated with 10 μM ruthenium red prior to and during hypotonic swelling (HS + RR) in order to inhibit all TRPV channels, while only TRPV4 was inhibited by treating ATDC5 cells with 10 μM RN1734 prior to and during hypotonic swelling (HS + RN). (* indicates p-value < 0.05 compared to controls)

### IGF-1 alters the mechanosensitivity through actin cytoskeletal polymerization

Increases in cell stiffness induced by actin cytoskeletal polymerization exerts a large influence on the sensitivity of cells to mechanical stimuli (Gardinier et al., 2014; Zhang et al., 2006). To determine if IGF-1 down-regulates channel activity during mechanical loading (Fig 1) by increasing ASFF (Fig 2), ATDC5 cells were pretreated with 300 ng/ml of IGF-1 for 3 hrs prior to application of mechanical stimulation with HS (Fig. 4A and B). IGF-1 pretreatment significantly reduced both the number of cells responding to HS and the magnitude of the [Ca^2+^]_i_ response to this stimulation. Addition of 1 μM of cytochalasin D five minutes before HS stimulation reversed the IGF-1-induced loss in mechanosensitivity, increasing both the number of responding cells and the magnitude of the [Ca^2+^]_i_ response compared to chondrocytes treated with IGF-1 alone. To determine the optimal dose of IGF-1that suppresses the TRPV4 channel, ATDC5 cells were treated with either 0, 1, 10, 100, or 300 ng/ml of IGF-1 for 3 hours prior to mechanical stimulation with HS. IGF-1 at a concentration of 100 ng/ml significantly reduced the magnitude of the [Ca^2+^]_i_ response (Fig 4C). To determine the optimal time of IGF-1 treatment, a time course of 0, 30, 60, 120 and 180 min was used when cells were treated with 300 ng/ml of IGF-1. A significant attenuation in the response to HS was achieved within 1 hour of treatment and sustained over a 3 hours of IGF-1 pretreatment (Fig. 4D).

**Figure 4:**
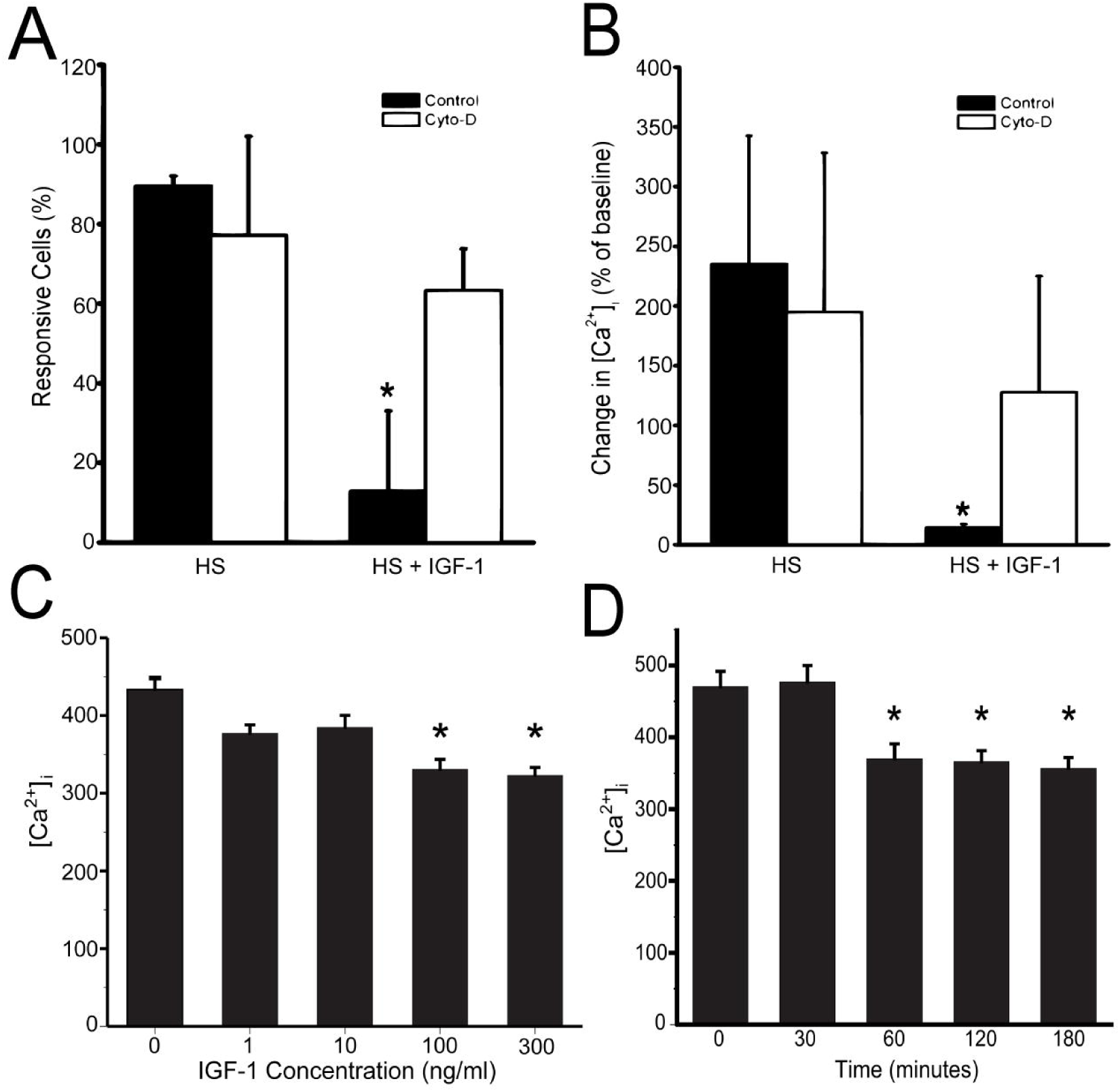
The [Ca^2+^]_i_ response to HS is altered by IGF-1 treatment and actin cytoskeleton organization. **A]** The number of cells responding to HS with a 50% or greater increase in [Ca^2+^]_i_ compared to baseline was significantly attenuated following IGF-1 treatment. **B]** The change in [Ca^2+^]_i_ compared to baseline in response to HS was significantly reduced following IGF-1 treatment. Disruption of the F-actin organization with 1 μM of cytochlasin D for 30 min after cells were treated with IGF-1 increased the number of cells exhibiting a [Ca^2+^]_i_ response to HS and magnitude in response. **C]** [Ca^2+^]_i_ of a dose response of ATDC5 to IGF-1 pre-treatment for 3 hr prior to challenge with 50% HS. Significant attenuation of [Ca^2+^]_I_ during hypotonic challenge occurs at concentrations of 100 ng/ml and 300 ng/ml of IGF-1. **D]** Attenuation of [Ca^2+^]_i_ when ATDC5 cells are challenged with HS starts within 60 minutes of IGF-1 treatment and is sustained over a 180 minute period.

We next sought to determine if the MAP7 binding domain of TRPV4 (aa. 798-809) is responsible for actin dependent regulation of TRPV4. HEK cells stably expressing WT and mutant (P799L and G800D) TRPV4 were pretreated with 300 ng/ml of IGF-1 for 3 hours prior to challenge with HS. IGF-1 pretreatment significantly attenuated the magnitude of [Ca^2+^]_i_ of the HEK-WT cells (Fig. 5). Untransfected HEK cells (UT) demonstrated a significantly lower [Ca^2+^]_i_ response to HS than wild type TRPV4 transfected (WT) cells when exposed to HS. Further UT cells exhibited no attenuation of the [Ca^2+^]_i_ signal during IGF-1 pretreatment. However, the [Ca^2+^]_i_ response of WT transfected HEK cells pretreated with IGF-1 was significantly reduced compared to WT controls indicating that TRPV4 transfection was mediating the increase in control WT cells. Both of the TRPV4 mutations, HEK-799 and HEK-800, responded to HS but neither cell line saw an attenuation in the [Ca^2+^]_i_ response to HS when pretreated with 300 ng/ml of IGF-1 for 3 hr. Interestingly, the HEK-800 cells were sensitized to HS when pretreated with IGF-1.

**Figure 5:**
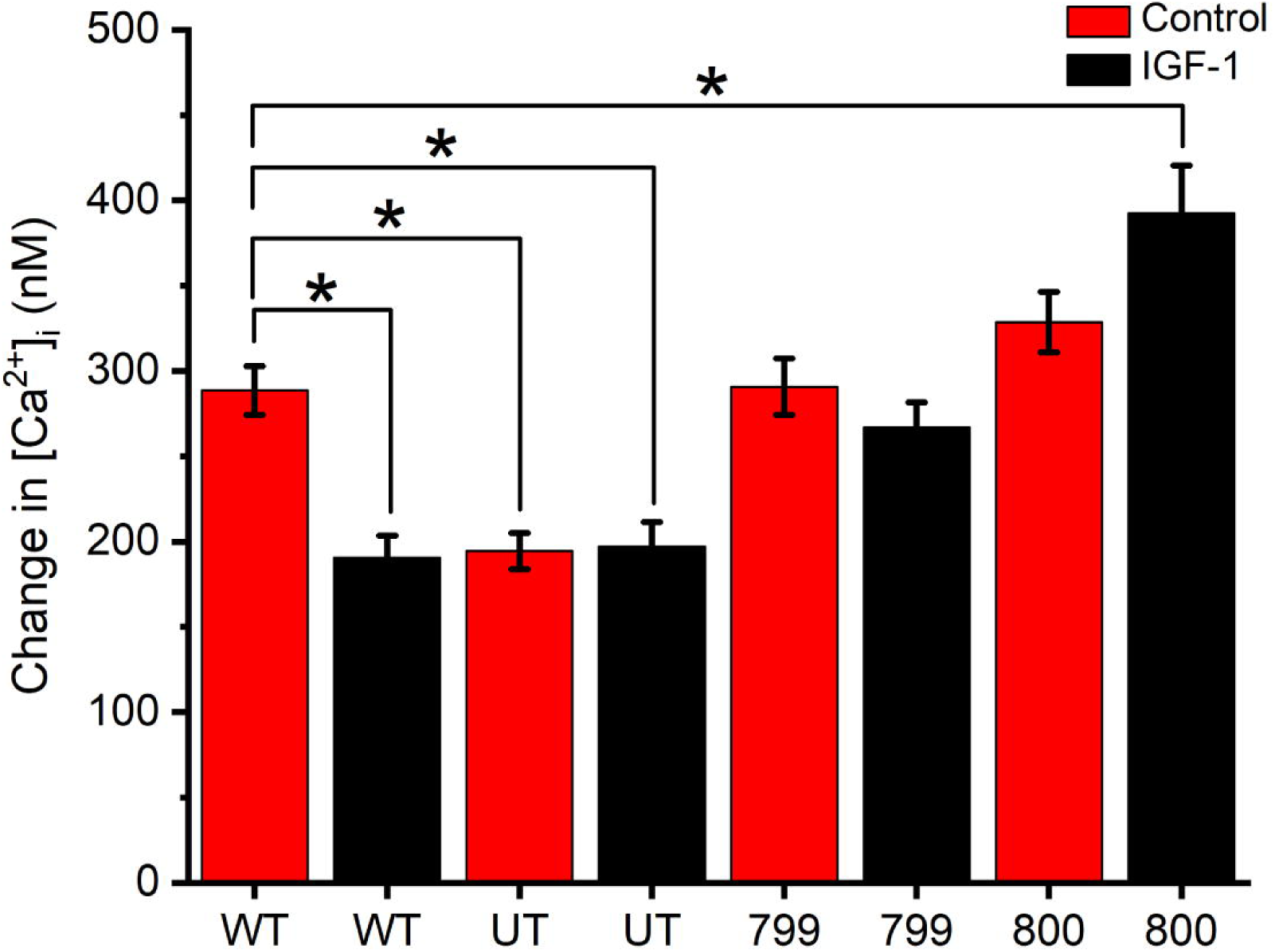
Effects of transfection of wild type TRPV4 or point mutations of the channel on calcium signaling in response to HS. The change in [Ca^2+^]_i_ compared to baseline in response to HS ± 300 ng/ml of IGF-1 in HEK 293 cells that were either un-transfected (UT) or stably transfected with either wild-type (WT) TRPV4, a mutation of proline to lysine at aa. 799 (799) TRPV4, or a mutation of glycine to aspartate at aa. 800 (800) TRPV4. IGF-1 treatment fails to suppress the response of these cells to HS in the 799 and 800 cells, indicating that actin interacts with TRPV4 within the MAP7 binding domain (aa. 798-809). (* indicates p-value < 0.05 compared across all groups)

### IGF-1 Alters Chondrocyte Purinergic Signaling

Purinergic signaling in response to mechanical loading is a vital component of the mechanotransduction pathway of chondrocytes (Millward-Sadler et al., 2004; Pingguan-Murphy et al., 2006). In response to HS, ATDC5 cells exhibited a significant release of ATP following 5 minutes exposure to HS (Fig 6A). This ATP release during HS was significantly attenuated by pretreatment with IGF-1 for 3 hours, but was fully restored when the actin cytoskeleton was disrupted using cytochalasin D. In addition, the general TRPV4 inhibitor, ruthenium red, or the TRPV4 specific inhibitor, RN1734, attenuated this response (Fig. 6B). These data suggest that ATP release during HS was mediated by TRPV channels, specifically TRPV4.

**Figure 6:**
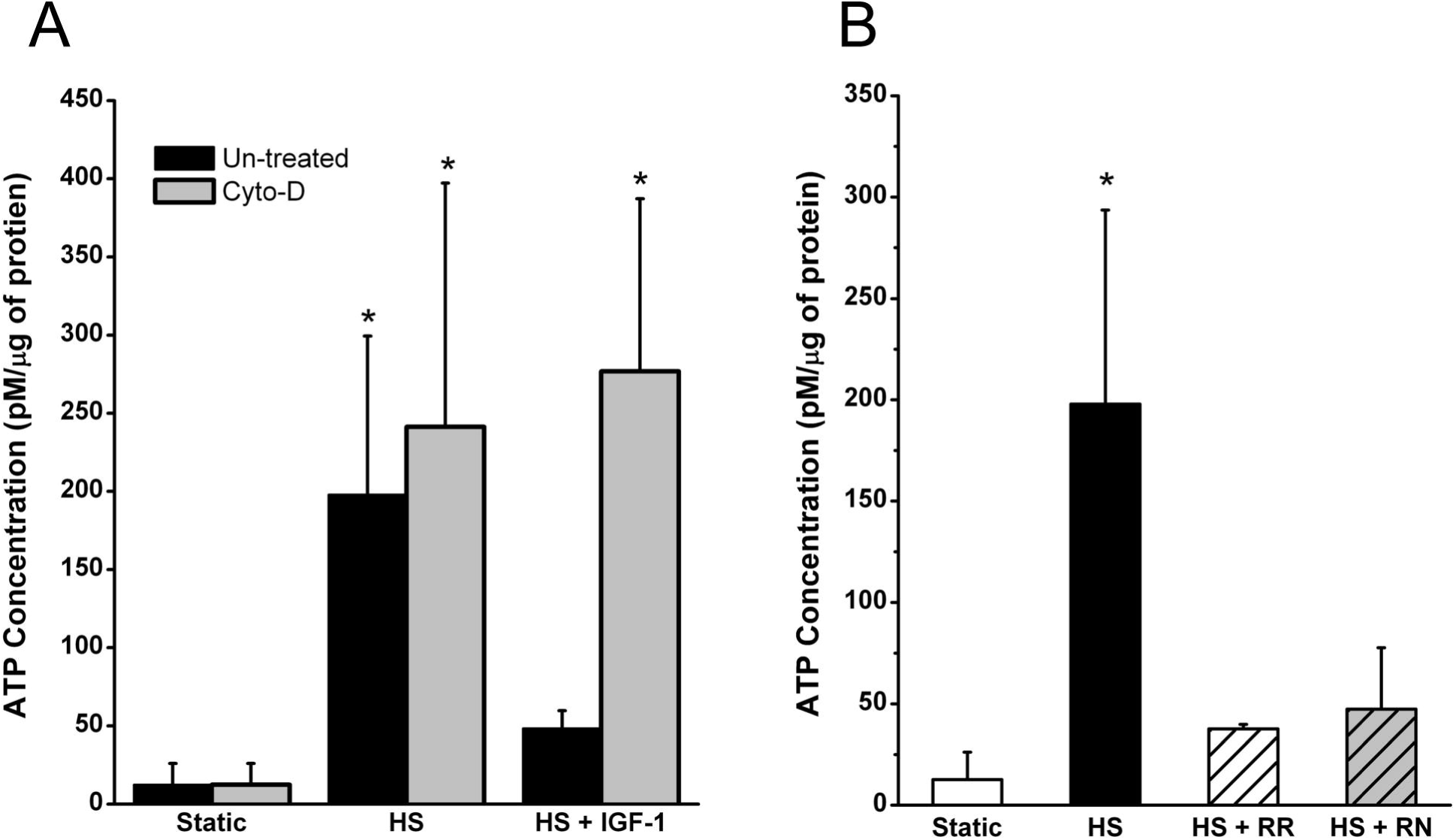
ATP release during HS is reduced following IGF-1 treatment and inhibition of TRPV channels. After 5 minutes of HS, ATP release was 12 times higher than static controls. IGF-1 treatment attenuated the release of ATP and was only 3 times greater than static controls. Inhibition of TRPV channels reduced ATP release during HS (HS + RR), and inhibition of TRPV4 (HS + RN) specifically reduced ATP by the same degree. (* indicates a p-value < 0.05 compared to controls)

## Discussion

This study elucidates a novel mechanism by which IGF-1 interacts with mechanical stimuli in regulating TRPV4 channel kinetics. Specifically, the data indicate that IGF-1 increases cell stiffness through regulation of the actin cytoskeleton and thereby suppress the activation of TRPV4 channels in response to mechanical stimulation. Several reports have suggested that actin and microtubules may be closely associated with this channel (Becker et al., 2009; Goswami et al., 2010; Suzuki et al., 2003). Perhaps the strongest indication that actin regulates this channel is that when actin is depolymerized, the osmotic response of the cell is lost (Becker et al., 2009; Suzuki et al., 2003). While it is unclear whether actin directly interacts with the TRPV4 channel, Suzuki, et al. have shown that the TRPV4 channel expresses the Microtubulin Associating Protein 7 (MAP7) domain at the amino acid sequence 798-809 of the channel and that actin, but not microtubules, regulates channel activity (Suzuki et al., 2003). These reports, coupled with the data shown here, suggest that IGF-1 can directly regulate TRPV4 channel activity by increasing actin polymerization to reduce mechanical gating of the channel. As we have previously shown, ATP release from osteoblasts is dependent on calcium entry into the cell (Genetos et al., 2005). Thus, reducing the Ca^2+^ entry through the TRPV4 channel by IGF-1 could also impact ATP release in response to mechanical loading in chondrocytes.

Osteoarthritis is characterized by the degeneration of the cartilage matrix, resulting in fibrillation, tears, and eventual loss of both matrix and cells. Articular cartilage is normally a reservoir for water, as proteoglycans and collagen glycosaminoglycans retain water in both bound and unbound states. Damage to the matrix collagen network enables fibrillated cartilage to increase its hydration through changes in the local osmotic pressure gradient (Maroudas et al. 1979). In patients with OA, articular cartilage water content is increased and cartilage swelling is one of the earliest macroscopic events in OA progression (Liess et al 2002, Maroudas 1976, Jones et al., 1999).

At the cellular level, it is possible that chondrocytes in articular cartilage experience an increase in cell volume in OA. It has been estimated that chondrocyte volume can increase up to ~90% in high-grade OA cartilage, leading to cell swelling and inability to deform the membrane (Bush et al., 2003). Primary human chondrocytes isolated from OA patients exhibit an inability to regulate their cell volume compared to chondrocytes isolated from patients without OA (Jones et al., 1999). Subsequently, loss of chondrocyte number significantly attenuates any repair or maintenance of cartilage as these cells are responsible for matrix production.

Loss of local matrix likely plays a role in reduced chondrocyte anabolic activity. IGF-1 is normally located in the local matrix around chondrocytes, where it is available at relatively constant levels and signals to the cell to increase anabolic activity and for production of matrix molecules, such as collagen II and aggrecan (Martin et al., 2000). Thus, the significant reduction in IGF-1 levels within arthritic cartilage may contribute to the associated increase in chondrocyte death by not allowing these cells to maintain their structural integrity and withstand hypotonic conditions. Furthermore, autologous implantation of chondrocytes overexpressing IGF-1 significantly improves collagen production, glycosaminoglycan, and mechanical properties of articular cartilage in an equine defect model of PTOA (Ortved et al., 2015, Griffin et al., 2016).

TRPV4 has also been implicated in OA. Recent studies show that knock-down of TRPV4 in mice results in severe OA that is sex and age dependent. Furthermore, chondrocytes from these mice lack a calcium response to hypotonic conditions (Clark et al., 2010). Our study demonstrates that the TRPV4 channel plays a vital role in mediating intracellular calcium mobilization and subsequently purinergic signaling in response to hypotonic conditions. Other studies have demonstrated similar findings when TRPV4 is blocked under hypotonic conditions, along with a delayed regulatory volume decrease, and increase in prostaglandin E_2_ production (Phan et al., 2009).

Our study demonstrates a unique link between TRPV4 activity and IGF-1 mediated changes in the actin cytoskeleton. While Lee et al. has shown that IGF-1 sensitized TRPV4 activity through SGK1 phosphorylation of the serine 824 amino acid, 100 μM of IGF-1 was utilized in these studies (Lee et al., 2010). This supraphysiologic amount of IGF-1 is over 1000-fold higher than the concentration used in our studies. While this level of IGF-1 could induce numerous signaling pathways independent of TRPV4, it could indicate that TRPV4 may have a biphasic response to IGF-1 treatment that is concentration dependent. Although the mechanism by which IGF-1 alters actin polymerization is relatively unclear, Novakofski, et al. has shown that IGF-1 suppresses RhoA GTPase activation in chondrocytes, while increasing cortical F-actin staining below the plasma membrane (Novakofski et al., 2009). The cytoskeleton is tightly regulated by the Rho family of small GTPases, in particular RhoA, Rac1 and cdc42, and their coordinated activation in which activation of RhoA, for example, suppresses the activity of Rac1 and vice versa (Moorman et al., 1999). This is especially true in chondrocytes where activation of Rac1/cdc42 reduced proliferation and increased gene expression and hypertrophy by restricting activation of RhoA (Wang et al., 2005). Thus, IGF-1 is thought to activate Rac1 and increase the organization of cortical actin, which in turn stiffens the plasma membrane. This increase in cell stiffness then reduces the deformation of the plasma membrane under hypotonic conditions and the subsequent activation of the TRPV4 channel. In this study, we show that mutations within the MAP7 binding domain of TRPV4 influences the response of TRPV4 to IGF-1 treatment. As Suzuki et al. suggest, this domain is responsible for actin binding to the TRPV4 channel. The differential response to IGF-1 pretreatment in HEK-WT, HEK-799, and HEK-800 cells highlights the importance of this binding domain in actin-mediated regulation of TRPV4. Similar connections have been established in HaCaT keratinocytes and CHO cells transfected with TRPV4, indicating a link between TRPV4, F-actin and the cellular response to hypotonic swelling (Becker et al., 2009).

In this study, we highlight the link between IGF-1 regulation of TRPV4 and how it influences extracellular ATP release from chondrocytes. We show that IGF-1 pretreatment significantly attenuates the release of ATP from chondrocytes by modifying TRPV4 function through the actin cytoskeleton While chondrocytes release ATP in response to mechanical stimuli to promote chondrogenesis, elevated levels of ATP have been found in the synovial fluid of patients with chondrocalcinosis and OA (Ryan et al., 1992). This indicates that differential responses to extracellular ATP in chondrocytes may be influenced by the pathological status of cartilage. With the loss of IGF-1 seen in OA patients, mechanical loading of chondrocytes would lead to excessive ATP release. Depletion of ATP in chondrocytes has been previously reported in spontaneous OA in Hartley Guinea Pigs (Johnson et al., 2004). This leads to altered ATP homeostasis, which is known to drive OA and can be inhibited with exogenous adenosine administration (Corciulo et al., 2015)

Our results indicate that the release of ATP by chondrocytes is dependent on TRPV4 mediated calcium mobilization, which has been noted in other tissues. With inhibition of the channel, less efflux of ATP occurs from chondrocytes. While IGF-1 maintains chondrocytes homeostasis and promotes a chondrogenic phenotype, inhibition or knockout of TRPV4 suppresses chondrogenic ECM protein production and the responsiveness of chondrocytes to the beneficial effects of dynamic mechanical load. Thus, biochemical modulation of TRPV4 by IGF-1 may impact chondrocytes mechanosensation to impart an anabolic response in cartilage.

IGF-1 is a promising candidate for the prevention and treatment of articular cartilage damage in OA. Our data elucidate a potentially important mechanism by which IGF-1 regulates chondrocyte-like cells mechanosensitivity, a major contributor to articular chondrocyte function. This mechanism involves the coordinated control of actin organization, cell stiffness, TRPV4 channel activation and calcium responses. However, because mechanical signals regulate both anabolic and catabolic chondrocyte functions, future studies are needed to define the relationship between this regulatory pathway and chondrocyte anabolic and catabolic activities.

## Acknowledgements

P30 grant (Binder MacLeod) and NSF grant vouchers (Steven)

## Notes

#### Summary of Updates

Addition of Authors

## REFERENCES

Becker, D., Bereiter-Hahn, J., Jendrach, M., 2009. Functional interaction of the cation channel transient receptor potential vanilloid 4 (TRPV4) and actin in volume regulation. Eur. J. Cell Biol. 88, 141–152. https://doi.org/10.1016/j.ejcb.2008.10.002

Bonassar, L.J., Grodzinsky, A.J., Frank, E.H., Davila, S.G., Bhaktav, N.R., Trippel, S.B., 2001. The effect of dynamic compression on the response of articular cartilage to insulin-like growth factor-I. J. Orthop. Res. 19, 11–17. https://doi.org/10.1016/S0736-0266(00)00004-8

Bonassar, L.J., Grodzinsky, A.J., Srinivasan, A., Davila, S.G., Trippel, S.B., 2000. Mechanical and Physicochemical Regulation of the Action of Insulin-Like Growth Factor-I on Articular Cartilage. Arch. Biochem. Biophys. 379, 57–63. https://doi.org/10.1006/ABBI.2000.1820

Bush, P.., Hall, A.., 2003. The volume and morphology of chondrocytes within non-degenerate and degenerate human articular cartilage. Osteoarthr. Cartil. 11, 242–251. https://doi.org/10.1016/S1063-4584(02)00369-2

Carter, D.R., Beaupr??, G.S., Wong, M., Smith, R.L., Andriacchi, T.P., Schurman, D.J., 2004. The Mechanobiology of Articular Cartilage Development and Degeneration. Clin. Orthop. Relat. Res. 427, S69–S77. https://doi.org/10.1097/01.blo.0000144970.05107.7e

Center for Disease Control. 2010. Osteoarthritis. http://www.cdc.gov/arthritis/basics/osteoarthritis.htm.

Clark, A.L., Votta, B.J., Kumar, S., Liedtke, W., Guilak, F., 2010. Chondroprotective role of the osmotically sensitive ion channel transient receptor potential vanilloid 4: Age+ and sex+ dependent progression of osteoarthritis in Trpv4-deficient mice. Arthritis Rheum. 62, 2973–2983. https://doi.org/10.1002/art.27624

Corciulo, C., Lendhey, M., Wilder, T., Schoen, H., Cornelissen, A.S., Chang, G., Kennedy, O.D., Cronstein, B.N., 2017. Endogenous adenosine maintains cartilage homeostasis and exogenous adenosine inhibits osteoarthritis progression. Nat. Commun. 8, 15019. https://doi.org/10.1038/ncomms15019

De Croos, J.N.A., Dhaliwal, S.S., Grynpas, M.D., Pilliar, R.M., Kandel, R.A., 2006. Cyclic compressive mechanical stimulation induces sequential catabolic and anabolic gene changes in chondrocytes resulting in increased extracellular matrix accumulation. Matrix Biol. 25, 323–331. https://doi.org/10.1016/J.MATBIO.2006.03.005

Ellsworth, J.L., Berry, J., Bukowski, T., Claus, J., Feldhaus, A., Holderman, S., Holdren, M.S., Lum, K.D., Moore, E.E., Raymond, F., Ren, H.P., Shea, P., Sprecher, C., Storey, H., Thompson, D.L., Waggie, K., Yao, L., Fernandes, R.J., Eyre, D.R., Hughes, S.D., 2002. Fibroblast growth factor-18 is a trophic factor for mature chondrocytes and their progenitors. Osteoarthr. Cartil. 10, 308–320. https://doi.org/10.1053/joca.2002.0514

Fanning, P.J., Emkey, G., Smith, R.J., Grodzinsky, A.J., Szasz, N., Trippel, S.B., 2003. Mechanical regulation of mitogen-activated protein kinase signaling in articular cartilage. J. Biol. Chem. 278, 50940–8. https://doi.org/10.1074/jbc.M305107200

Flockerzi, V., Nilius, B. (Eds.), 2007. Transient Receptor Potential (TRP) Channels, Handbook of Experimental Pharmacology. Springer Berlin Heidelberg, Berlin, Heidelberg. https://doi.org/10.1007/978-3-540-34891-7

Fortier, L.A., Deak, M.M., Semevolos, S.A., Cerione, R.A., 2004. Insulin-like growth factor-I diminishes the activation status and expression of the small GTPase Cdc42 in articular chondrocytes. J. Orthop. Res. 22, 436–445. https://doi.org/10.1016/j.orthres.2003.08.021

Gao, X., Wu, L., O’Neil, R.G., 2003. Temperature-modulated diversity of TRPV4 channel gating: activation by physical stresses and phorbol ester derivatives through protein kinase C-dependent and -independent pathways. J. Biol. Chem. 278, 27129–37. https://doi.org/10.1074/jbc.M302517200

Gardinier, J., Yang, W., Madden, G.R., Kronbergs, A., Gangadharan, V., Adams, E., Czymmek, K., Duncan, R.L., 2014. P2Y _2_ receptors regulate osteoblast mechanosensitivity during fluid flow. Am. J. Physiol. Physiol. 306, C1058–C1067. https://doi.org/10.1152/ajpcell.00254.2013

Goswami, C., Kuhn, J., Heppenstall, P.A., Hucho, T., 2010. Importance of non-selective cation channel TRPV4 interaction with cytoskeleton and their reciprocal regulations in cultured cells. PLoS One 5, 19–21. https://doi.org/10.1371/journal.pone.0011654

Griffin, D.J., Ortved, K.F., Nixon, A.J., Bonassar, L.J., 2016. Mechanical properties and structure-function relationships in articular cartilage repaired using IGF-I gene-enhanced chondrocytes. J. Orthop. Res. 34, 149–153. https://doi.org/10.1002/jor.23038

Haudenschild, D.R., Nguyen, B., Chen, J., D’Lima, D.D., Lotz, M.K., 2008. Rho kinase-dependent CCL20 induced by dynamic compression of human chondrocytes. Arthritis Rheum. 58, 2735–42. https://doi.org/10.1002/art.23797

Helmick, C.G., Felson, D.T., Lawrence, R.C., Gabriel, S., Hirsch, R., Kwoh, C.K., Liang, M.H., Kremers, H.M., Mayes, M.D., Merkel, P.A., Pillemer, S.R., Reveille, J.D., Stone, J.H., 2008. Estimates of the prevalence of arthritis and other rheumatic conditions in the United States: Part I. Arthritis Rheum. 58, 15–25. https://doi.org/10.1002/art.23177

Hurd, L., Kirwin, S.M., Boggs, M., Mackenzie, W.G., Bober, M.B., Funanage, V.L., Duncan, R.L., 2015. A mutation in TRPV4 results in altered chondrocyte calcium signaling in severe metatropic dysplasia. Am. J. Med. Genet. Part A 167, 2286–2293. https://doi.org/10.1002/ajmg.a.37182

Jin, M., Emkey, G.R., Siparsky, P., Trippel, S.B., Grodzinsky, A.J., 2003. Combined effects of dynamic tissue shear deformation and insulin-like growth factor I on chondrocyte biosynthesis in cartilage explants. Arch. Biochem. Biophys. 414, 223–31. https://doi.org/10.1016/s0003-9861(03)00195-4

Johnson, K., Svensson, C.I., Etten, D. Van, Ghosh, S.S., Murphy, A.N., Powell, H.C., Terkeltaub, R., 2004. Mediation of spontaneous knee osteoarthritis by progressive chondrocyte ATP depletion in Hartley guinea pigs. Arthritis Rheum. 50, 1216–1225. https://doi.org/10.1002/art.20149

Jones, W.R., Ping Ting-Beall, H., Lee, G.M., Kelley, S.S., Hochmuth, R.M., Guilak, F., 1999. Alterations in the Young’s modulus and volumetric properties of chondrocytes isolated from normal and osteoarthritic human cartilage. J. Biomech. 32, 119–127. https://doi.org/10.1016/S0021-9290(98)00166-3

Kim, Y.J., Sah, R.L.Y., Grodzinsky, A.J., Plaas, A.H.K., Sandy, J.D., 1994. Mechanical regulation of cartilage biosynthetic behavior: Physical stimuli. Arch. Biochem. Biophys. https://doi.org/10.1006/abbi.1994.1201

Lee, E.J., Shin, S.H., Chun, J., Hyun, S., Kim, Y., Kang, S.S., 2010. The modulation of TRPV4 channel activity through its Ser 824 residue phosphorylation by SGK1. Animal Cells Syst. (Seoul). 14, 99–114. https://doi.org/10.1080/19768354.2010.486939

Lee, H.-S., Millward-Sadler, S.J., Wright, M.O., Nuki, G., Al-Jamal, R., Salter, D.M., 2002. Activation of Integrin—RACK1/PKCα signalling in human articular chondrocyte mechanotransduction. Osteoarthr. Cartil. 10, 890–897. https://doi.org/10.1053/JOCA.2002.0842

Lee, H.S., Millward-Sadler, S.J., Wright, M.O., Nuki, G., Salter, D.M., 2000. Integrin and Mechanosensitive Ion Channel-Dependent Tyrosine Phosphorylation of Focal Adhesion Proteins and β-Catenin in Human Articular Chondrocytes After Mechanical Stimulation. J. Bone Miner. Res. 15, 1501–1509. https://doi.org/10.1359/jbmr.2000.15.8.1501

Leipzig, N.D., Eleswarapu, S.V., Athanasiou, K.A., 2006. The effects of TGF-β1 and IGF-I on the biomechanics and cytoskeleton of single chondrocytes. Osteoarthr. Cartil. 14, 1227–1236. https://doi.org/10.1016/J.JOCA.2006.05.013

Levin, S.M., 2000. Alterations in the Young’s modulus and volumetric properties of chondrocytes isolated from normal and osteoarthritic human cartilage (multiple letters). J. Biomech. https://doi.org/10.1016/S0021-9290(99)00147-5

Liedtke, W., 2006. Transient receptor potential vanilloid channels functioning in transduction of osmotic stimuli. J. Endocrinol. 191, 515–523. https://doi.org/10.1677/joe.1.07000

Liess, C., Lüsse, S., Karger, N., Heller, M., Glüer, C.-C., 2002. Detection of changes in cartilage water content using MRI T2-mapping in vivo. Osteoarthr. Cartil. 10, 907–913. https://doi.org/10.1053/joca.2002.0847

Loukin, S., Zhou, X., Su, Z., Saimi, Y., Kung, C., 2010. Wild-type and brachyolmia-causing mutant TRPV4 channels respond directly to stretch force. J. Biol. Chem. 285, 27176–81. https://doi.org/10.1074/jbc.M110.143370

Madry, H., Zurakowski, D., Trippel, S., 2001. Overexpression of human insulin-like growth factor-I promotes new tissue formation in an ex vivo model of articular chondrocyte transplantation. Gene Ther. 8, 1443–1449. https://doi.org/10.1038/sj.gt.3301535

Mak, A.F., Lai, W.M., Mow, V.C., 1987. Biphasic indentation of articular cartilage—I. Theoretical analysis. J. Biomech. 20, 703–714. https://doi.org/10.1016/0021-9290(87)90036-4

Mariani, E., Pulsatelli, L., Facchini, A., 2014. Signaling pathways in cartilage repair. Int. J. Mol. Sci. 15, 8667–98. https://doi.org/10.3390/ijms15058667

Maroudas, A., Venn, M., 1977. Chemical composition and swelling of normal and osteoarthritic femoral head cartilage. II. Swelling. Ann. Rheum. Dis. 36, 399–406. https://doi.org/10.1136/ard.36.5.399

Martin, J.A., Buckwalter, J.A., 2000. The role of chondrocyte-matrix interactions in maintaining and repairing articular cartilage. Biorheology 37, 129–40.

Matta, C., 2012. Calcium signalling in chondrogenesis implications for cartilage repair. Front. Biosci. S5, 305–324. https://doi.org/10.2741/s374

Matta, C., Fodor, J., Szíjgyártó, Z., Juhász, T., Gergely, P., Csernoch, L., Zákány, R., 2008. Cytosolic free Ca2+ concentration exhibits a characteristic temporal pattern during in vitro cartilage differentiation: A possible regulatory role of calcineurin in Ca-signalling of chondrogenic cells. Cell Calcium 44, 310–323. https://doi.org/10.1016/J.CECA.2007.12.010

Mawatari, T., Lindsey, D.P., Harris, A.H.S., Goodman, S.B., Maloney, W.J., Smith, R.L., 2010. Effects of tensile strain and fluid flow on osteoarthritic human chondrocyte metabolism in vitro. J. Orthop. Res. 28, n/a–n/a. https://doi.org/10.1002/jor.21085

Millward-Sadler, S.J., Wright, M.O., Flatman, P.W., Salter, D.M., 2004. ATP in the mechanotransduction pathway of normal human chondrocytes, Biorheology. IOS Press.

Mizoguchi, F., Mizuno, A., Hayata, T., Nakashima, K., Heller, S., Ushida, T., Sokabe, M., Miyasaka, N., Suzuki, M., Ezura, Y., Noda, M., 2008. Transient receptor potential vanilloid 4 deficiency suppresses unloading-induced bone loss. J. Cell. Physiol. 216, 47–53. https://doi.org/10.1002/jcp.21374

Moorman, J.P., Luu, D., Wickham, J., Bobak, D.A., Hahn, C.S., 1999. A balance of signaling by Rho family small GTPases RhoA, Rac1 and Cdc42 coordinates cytoskeletal morphology but not cell survival. Oncogene 18, 47–57. https://doi.org/10.1038/sj.onc.1202262

Muramatsu, S., Wakabayashi, M., Ohno, T., Amano, K., Ooishi, R., Sugahara, T., Shiojiri, S., Tashiro, K., Suzuki, Y., Nishimura, R., Kuhara, S., Sugano, S., Yoneda, T., Matsuda, A., 2007. Functional gene screening system identified TRPV4 as a regulator of chondrogenic differentiation. J. Biol. Chem. 282, 32158–32167. https://doi.org/10.1074/jbc.M706158200

Nebelung, S., Gavenis, K., Lüring, C., Zhou, B., Mueller-Rath, R., Stoffel, M., Tingart, M., Rath, B., 2012. Simultaneous anabolic and catabolic responses of human chondrocytes seeded in collagen hydrogels to long-term continuous dynamic compression. Ann. Anat. - Anat. Anzeiger 194, 351–358. https://doi.org/10.1016/J.AANAT.2011.12.008

Neu, C.P., Khalafi, A., Komvopoulos, K., Schmid, T.M., Reddi, A.H., 2007. Mechanotransduction of bovine articular cartilage superficial zone protein by transforming growth factor β signaling. Arthritis Rheum. 56, 3706–3714. https://doi.org/10.1002/art.23024

Nilius, B., 2004. TRPV4 calcium entry channel: a paradigm for gating diversity. AJP Cell Physiol. 286, 195C – 205. https://doi.org/10.1152/ajpcell.00365.2003

Nilius, B., Prenen, J., Wissenbach, U., Bödding, M., Droogmans, G., 2001. Differential activation of the volume-sensitive cation channel TRP12 (OTRPC4) and volume-regulated anion currents in HEK-293 cells. Pflugers Arch. Eur. J. Physiol. 443, 227–233. https://doi.org/10.1007/s004240100676

Nilius, B., Watanabe, H., Vriens, J., 2003. The TRPV4 channel: structure-function relationship and promiscuous gating behaviour. Pflügers Arch. - Eur. J. Physiol. 446, 298–303. https://doi.org/10.1007/s00424-003-1028-9

Novakofski, K., Boehm, A., Fortier, L., 2009. The small GTPase Rho mediates articular chondrocyte phenotype and morphology in response to interleukin-1α and insulin-like growth factor-I. J. Orthop. Res. 27, 58–64. https://doi.org/10.1002/jor.20717

Oh, C.-D., Chun, J.-S., 2003. Signaling mechanisms leading to the regulation of differentiation and apoptosis of articular chondrocytes by insulin-like growth factor-1. J. Biol. Chem. 278, 36563–71. https://doi.org/10.1074/jbc.M304857200

Ortved, K.F., Begum, L., Mohammed, H.O., Nixon, A.J., 2015. Implantation of rAAV5-IGF-I transduced autologous chondrocytes improves cartilage repair in full-thickness defects in the equine model. Mol. Ther. 23, 363–373. https://doi.org/10.1038/mt.2014.198

Pedersen, S.F., Owsianik, G., Nilius, B., 2005. TRP channels: An overview. Cell Calcium 38, 233–252. https://doi.org/10.1016/J.CECA.2005.06.028

Phan, M.N., Leddy, H.A., Votta, B.J., Kumar, S., Levy, D.S., Lipshutz, D.B., Lee, S.H., Liedtke, W., Guilak, F., 2009. Functional characterization of TRPV4 as an osmotically sensitive ion channel in porcine articular chondrocytes. Arthritis Rheum. 60, 3028–37. https://doi.org/10.1002/art.24799

Pingguan-Murphy, B., El-Azzeh, M., Bader, D.L., Knight, M.M., 2006. Cyclic compression of chondrocytes modulates a purinergic calcium signalling pathway in a strain rate+ and frequency-dependent manner. J. Cell. Physiol. 209, 389–397. https://doi.org/10.1002/jcp.20747

Ryan, L.M., Kurup, I. V., Derfus, B.A., Kushnaryov, V.M., 1992. ATPLJinduced chondrocalcinosis. Arthritis Rheum. 35, 1520–1525. https://doi.org/10.1002/art.1780351216

Sah, R.L.-Y., Doong, J.-Y.H., Grodzinsky, A.J., Plaas, A.H.K., Sandy, J.D., 1991. Effects of compression on the loss of newly synthesized proteoglycans and proteins from cartilage explants. Arch. Biochem. Biophys. 286, 20–29. https://doi.org/10.1016/0003-9861(91)90004-3

Suzuki, M., Hirao, A., Mizuno, A., 2003. Microtubule-associated [corrected] protein 7 increases the membrane expression of transient receptor potential vanilloid 4 (TRPV4). J. Biol. Chem. 278, 51448–53. https://doi.org/10.1074/jbc.M308212200

Trippel, S.B., 1995. Growth factor actions on articular cartilage. J. Rheumatol. Suppl. 43, 129–32.

Wang, G., Beier, F., 2005. Rac1/Cdc42 and RhoA GTPases Antagonistically Regulate Chondrocyte Proliferation, Hypertrophy, and Apoptosis. J. Bone Miner. Res. 20, 1022–1031. https://doi.org/10.1359/JBMR.050113

Wang, P., Zhu, F., Lee, N.H., Konstantopoulos, K., 2010. Shear-induced interleukin-6 synthesis in chondrocytes: roles of E prostanoid (EP) 2 and EP3 in cAMP/protein kinase A+ and PI3-K/Akt-dependent NF-kappaB activation. J. Biol. Chem. 285, 24793–804. https://doi.org/10.1074/jbc.M110.110320

Zhang, J., Ryder, K.D., Bethel, J.A., Ramirez, R., Duncan, R.L., 2006. PTH-Induced Actin Depolymerization Increases Mechanosensitive Channel Activity to Enhance Mechanically Stimulated 2+ Signaling in Osteoblasts*. J. Bone Miner. Res. 21, 1729–1737. https://doi.org/10.1359/jbmr.060722

